# Protamine lacunae preserve the paternal chromatin landscape in sperm

**DOI:** 10.1101/2025.10.03.680364

**Authors:** Thomas W Tullius, Rachel A Heuer, Stephanie C. Bohaczuk, Ben Mallory, Danilo Dubocanin, Jane Ranchalis, Ahmet Ayaz, Christopher E Mason, Emre Seli, Adam M Phillippy, Andrew B Stergachis, Bluma J Lesch

## Abstract

The transmission of the paternal genome requires extensive chromatin reorganization, in which nucleosomes are largely replaced by protamines that drive extreme condensation of the genome in the sperm head. Using Fiber-seq, we resolve patterns of paternal chromatin repackaging in sperm at single-molecule resolution, revealing the interplay between protamination and nucleosome retention. We find that nucleosome retention is probabilistic, with no locus universally occupied. Although promoters of spermatogenic genes preferentially harbor retained nucleosomes, the predominant carrier of paternal epigenetic information is protamine lacunae, accessible discontinuities in the protamine coat that preferentially mark critical regulatory elements. By contrast, centromere kinetochore binding regions robustly retain CENP-A mono-nucleosomes, providing a mechanism for the focal transmission of paternal centromeres. Finally, we find that paternal chromatin repackaging is altered in low-motility sperm. Together, these findings reveal distinct modes of paternal chromatin epigenetic inheritance with broad implications for development and infertility.

## Introduction

Sperm are essential for transmission of the paternal genome across generations. Faithful transmission requires pervasive repackaging of the genome, including eviction of most nucleosomes, coating of DNA with small basic protamine proteins, and reorganization of the genome into dense toroidal structures optimized for transit and fertilization ^1,2^. Despite this extensive reorganization, mature human sperm consistently retain 4-15% of their nucleosomes ^3–6^, hinting at potential roles for retained nucleosomes in early development or transgenerational epigenetic inheritance. However, whether nucleosomes are preferentially retained at specific genomic loci, and whether this retention holds functional significance for fertility or embryogenesis, remains unresolved.

Microscopy, *in situ* hybridization, and immunofluorescence studies have pointed to a stereotyped higher-order organization of chromosomes within human and murine sperm ^7,8^, and indicated enriched retention of nucleosomes in select locations, including the centromere-specific histone variant CENP-A at centromeric regions ^9–11^. ChIP-seq studies have made progress toward comprehensively mapping these retained nucleosomes ^5,6,12–19^, but have been confounded by contamination biases from rare somatic cells, amplification biases owing to the rarity of these retained nucleosomes, mapping biases of short reads to repetitive regions of the genome, and most fundamentally the inability to co-measure nucleosomes and protamines at the level of individual sperm. Consequently, our understanding of the landscape of nucleosome retention in sperm is largely incomplete. For example, while some studies show enrichment of nucleosomes at centromeric and repetitive elements ^13,14^, others report enrichment primarily at promoters of developmental genes ^5,6,12,18,20^.

Here, we demonstrate that Fiber-seq can accurately map nucleosome retention and protamine footprints simultaneously from telomere to telomere at the level of individual sperm. Fiber-seq is a long-read, single-molecule chromatin fiber sequencing method that uses a non-specific adenine methyltransferase to stencil protein footprints along underlying chromatin fibers with near nucleotide resolution ^21^. We validate that Fiber-seq can accurately distinguish protamine and nucleosome footprints in sperm and demonstrate that non-random nucleosome retention does indeed occur in human sperm, with the majority of retained nucleosomes existing as mono- and di-nucleosomes within patches of protamine-depleted DNA and surrounded by extended regions of protamination. Application of Fiber-seq to *in vitro* reconstituted protaminated DNA, normal and abnormal sperm, and human testes exposes that nucleosome retention and protamination are modulated by sequence and disease-state dependencies, with sperm retaining non-genetic information via two main modes: (1) strong retention of nucleosomes, exemplified by CENP-A at centromeric regions; or (2) more commonly by systematic depletion of protamines (protamine lacunae), exemplified by sites of high transcription during late sperm development. Additionally, we find that excessive protamine replacement and failure to appropriately retain nucleosomes are associated with male fertility defects, and retention of CENP-A footprints at centromere dip regions (CDRs) supports the inheritance of focal kinetochore attachment points within the human centromere.

### Single-molecule mapping of protamine-DNA footprints

We first sought to determine whether we could uniquely distinguish the DNA occupancy patterns of protamines using long-read single-molecule stencilling with Fiber-seq (**Fig. 1A**). Prior *in vitro* work has demonstrated that protamines condense DNA into toroids containing tens of kilobases of DNA, distinct from the typical ∼146bp stretches occupied by histones ^2,22–24^. To evaluate detection of this state by Fiber-seq, we reconstituted protaminated DNA *in vitro* by incubating human genomic DNA with increasing concentrations of purified human protamine proteins. The resultant protamine-DNA complexes were then treated with the non-specific m6A-methyltransferase Hia5 to stencil regions of protein occupancy. Overall, we observed that high protamine concentrations were associated with low m6A methylation (**Fig. 1B**), indicating that protamines occupy DNA in a dose-dependent manner that occludes the underlying DNA from m6A methylation.

**Figure 1.**
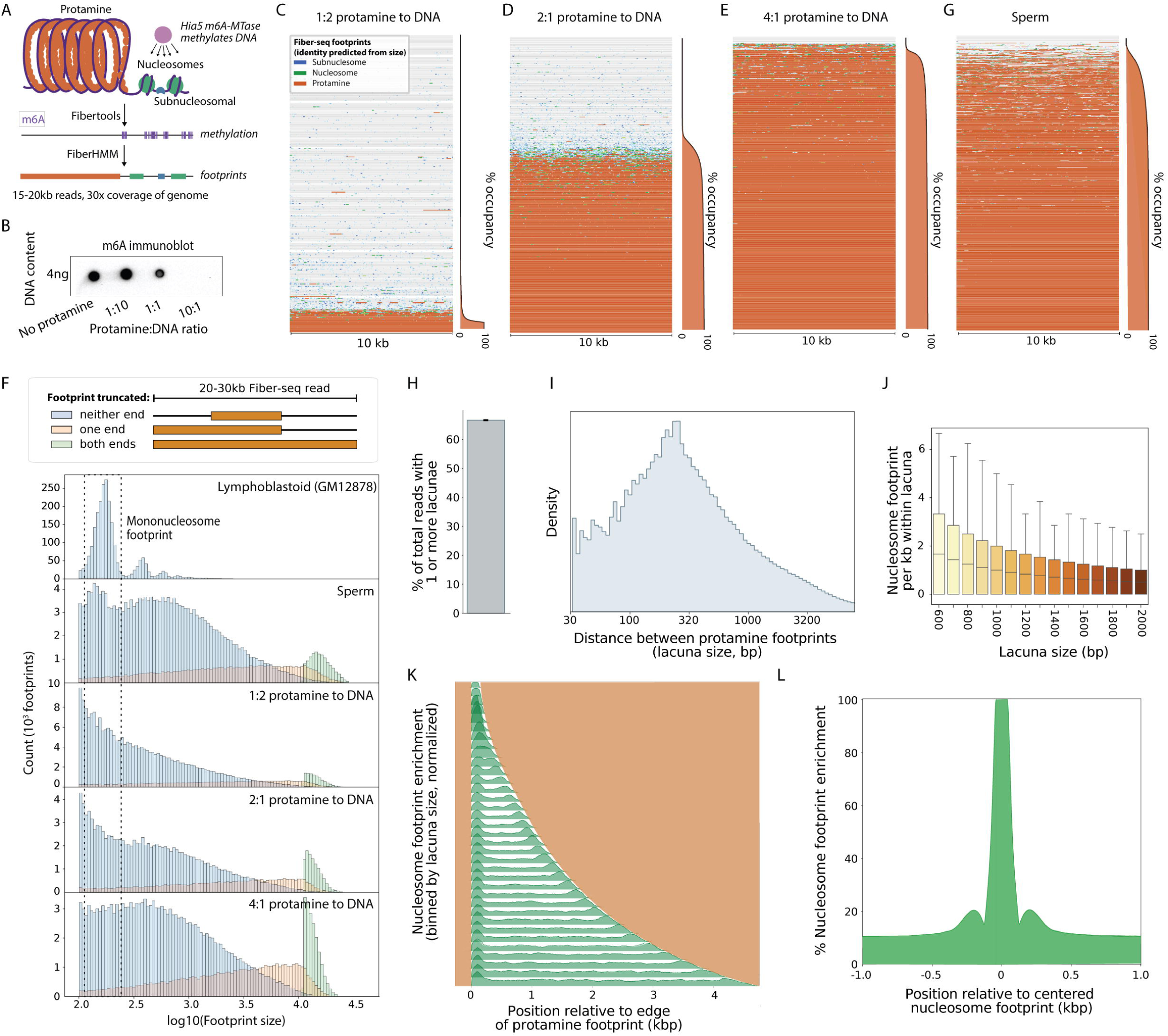
Sperm protamine profiles called by Fiber-seq. **A.** Schematic of sperm chromatin showing protamines (orange) and nucleosomes (green) protecting DNA from nonspecific m6A methylation by Hia5 (top), methylation footprinting (middle), and subsequent calling of protamine, nucleosomal, and subnucleosomal footprints (bottom). **B.** Dot blot for m6A in human genomic DNA reconstituted with varying ratios of DNA:human protamine mixture *in vitro*. **C** to **E.** Individual Fiber-seq reads (n = 500) from DNA reconstituted with protamines at the ratios indicated. Reads are sorted from low to high protamine footprint coverage content and a summary of percent protamine occupancy for each read is shown at right. Orange, protamine footprints; green, nucleosomal footprints; blue, subnucleosomal footprints. **F.** Distributions of footprint sizes in reconstituted samples at different protamine:DNA ratios, compared to human sperm and lymphoblastoid cells (from n = 10^6^ sampled reads). Separate distributions are shown for reads where protamine blocks are not truncated (blue), truncated at one end (orange), or truncated at both ends (green, indicating that the entire read is protected by a protamine block). **G.** Individual Fiber-seq reads from human sperm chromatin sorted from low to high protamine content as in panels C-E. **H.** Percentage of total sperm reads (from n = 10^6^ sampled reads) containing discontinuous protamine blocks (protamine lacunae). Errorbar indicates 95% confidence interval from 1000x bootstrapped resamplings. **I.** Histogram of lacuna size (distance between adjacent protamine footprints) in sperm (from n = 10^6^ sampled reads). **J.** Numbers of nucleosomal footprints called within lacunae of different sizes. Only lacunae long enough to theoretically accommodate multiple nucleosomes (>600 bp) are shown (from n = 10^6^ sampled reads). **K.** Metagenes of nucleosome position relative to the edge of a protamine footprint, binned by lacuna size (from n = 10^6^ sampled reads). **L.** Metagene showing distance from a nucleosome footprint (centered) to the nearest nucleosome footprint (from n = 10^6^ sampled reads).

To determine the architecture of these protamine footprints, we sequenced each sample using long-read PacBio HiFi sequencing to identify regions along each DNA molecule that were protected from m6A methylation (*i.e.,* single-molecule protein occupancy footprints) (**Fig. 1A, Fig. S1A-C**). Application of FiberHMM ^25^ to these sequencing reads demonstrated that *in vitro* protaminated DNA preferentially harbors large >1 kb protein footprints in a dose-dependent manner (**Fig. 1C-F, Supplemental Fig. S1A-C**). Notably, we observed that footprint patterns markedly differed between individual chromatin fibers, where a given molecule often displayed either near-complete protection from the m6A-MTase, or near-complete methylation (**Fig. 1C-E; Supplemental Fig. S1A-C**). Partially protected fibers were markedly enriched for having small (<100 bp) footprints, likely representing small separated protamine blocks, within the transition region between >1kb footprints and neighboring accessible regions (**Fig. 1F, Supplemental Fig. S1B**). Overall, these *in vitro* findings demonstrate that while protamines are capable of forming short DNA occupancy footprints, protamines most often co-bind DNA in a manner that results in large >1kb footprints with a distinct Fiber-seq signature, consistent with cooperative assembly of protamine-DNA complexes ^26,27^.

### Mono-nucleosomes and protamines co-occupy sperm chromatin fibers

Having established that protamines preferentially form large multi-kilobase m6A-MTase footprints *in vitro*, we next wanted to determine whether sperm display similar footprint patterns. We purified human ejaculated sperm via density gradient to ensure minimal contamination from non-sperm cells, and subjected these samples to Fiber-seq using a protocol optimized for sperm. m6A-MTase stenciled sperm fibers were then sequenced using PacBio single-molecule sequencing, and m6A-MTase protected footprints were identified using FiberHMM. Notably, the single-molecule chromatin architectures obtained by applying Fiber-seq to sperm markedly diverged from those obtained from other human cell lines and tissues. Specifically, whereas somatic tissue chromatin stencils largely contain regularly spaced ∼146 bp protected nucleosomal footprints bookended by ∼60 bp m6A-MTase accessible internucleosomal linker regions ^21,25^, sperm chromatin stencils are dominated by long (>1kb) footprints bookended by m6A-MTase accessible regions of varying size, a pattern consistent with that observed in the protamine reconstitution experiments (**Fig. 1F-G**, **Supplemental Fig. S1B-E)**. However, unlike the *in vitro* protamine reconstitution stencils where nucleosomes were not present, sperm chromatin stencils also contained a preponderance of ∼146bp footprints, indicating that Fiber-seq can accurately delineate nucleosome-sized footprints from protamine-sized footprints with single-molecule and near nucleotide resolution (**Fig. 1F-G**, **Supplemental Fig. S1B-E)**.

Using these single-molecule stencils, we next evaluated how protamine and nucleosome footprints are co-distributed in normal human sperm. As our sequencing read lengths were only 15-20kb on average, we were unable to quantify the maximum size of a protamine block in sperm. Overall, 33% of sperm fibers contained stencils consistent with being entirely protected by a large protamine footprint, indicating that a significant minority of protamine blocks are greater than 15kb in length (**Fig. 1H**). However, the majority (67%) of sperm fibers contained at least one patch of highly accessible DNA intermingled within extended protamine footprints, suggesting that a substantial fraction of sperm DNA is packaged into intermediate-length protamine blocks interrupted by short protamine gaps (*i.e.,* protamine lacunae) (**Fig. 1H-I**). Most protamine lacunae were short (<1,000 base pairs) and contained 0 or 1 nucleosome-sized footprints (**Figure 1I-J**). Notably, even when protamine lacunae were long enough to accommodate multiple nucleosomes, the majority contained only a single nucleosome or less commonly a pair of adjacent nucleosomes (**Fig. 1J**). Nucleosomes were enriched at the edges of protamine lacunae, supporting a potential role for stable nucleosomes as barriers to protamine domains (**Fig. 1K**). When paired nucleosomes were present within a protamine lacuna, their positioning relative to each other was similar to that observed within somatic chromatinized DNA (**Fig. 1L, Supplemental Fig. S1F**), suggesting that these nucleosome pairs may have been co-retained during spermiogenesis.

### Mapping the nucleosome-to-protamine transition in testis

Having observed that nucleosomes are present within protamine lacunae, we sought to understand how nucleosomes are selected for retention. To evaluate the process of nucleosome eviction and protamination during the human histone-to-protamine transition, we performed Fiber-seq on whole human testis tissue, as well as other somatic tissues from the same human donor as a comparison ^28^. Because protaminated sperm are continually produced from normally chromatinized spermatogenic stem cells throughout adulthood, adult testis contain spermatogenic cells at all stages of nucleosome eviction and protamination, starting from fully nucleosomal, chromatinized precursors and ending at protaminated sperm ready to enter the epididymis (**Fig. 2A**). We observed that whereas Fiber-seq data from non-testis tissues was predominantly marked by ∼146 bp footprints, testis Fiber-seq data showed both ∼146 bp footprints as well as larger >2 kb footprints (**Fig. 2B-C, Supplemental Fig. S2A-B**), consistent with the presence of protaminated chromatin fibers uniquely within the testis tissue.

**Figure 2.**
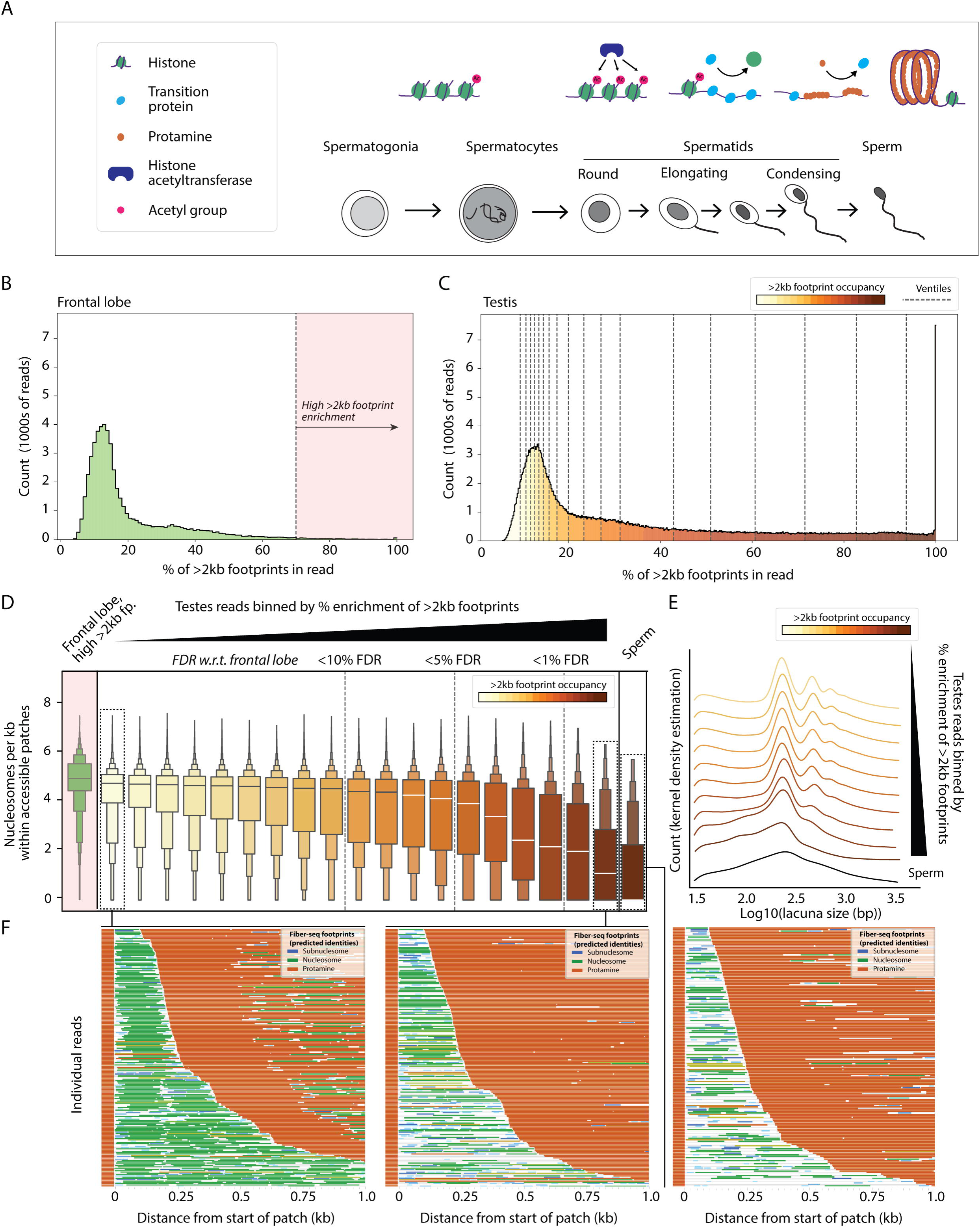
Nucleosomes are progressively evicted from accessible patches during protamine deposition. **A.** Schematic of spermatogenic development showing progression of histone-to-protamine exchange during post-meiotic spermatid development, including histone hyperacetylation, transition protein loading, and protamine deposition. **B.** Histogram showing % occupancy of reads by >2 kilobase footprints in Fiber-seq reads derived from the frontal lobe (n = 56,368). Dashed line and arrow indicates 70% cutoff used for high >2kb footprint reads used for boxplots in Figure 2D. **C.** Histogram showing % occupancy of reads by >2 kilobase footprints in Fiber-seq reads derived from testis (n = 9,149,319). Dashed lines and color gradient indicate ventile bins used for boxplots in Figure 2D. **D.** Nucleosome density in accessible patches in ventile bins of whole-testis Fiber-seq reads sorted by enrichment of >2 kb protamine-like footprints. Color gradient represents enrichment from Figure 1C. Nucleosome density within lacunae (n = 371,430, right, gradient boxenplot) in sperm (right boxplots) and frontal lobe (n = 78, left, green boxenplot) are shown for comparison. Frontal lobe plot shows all reads (n = 1432) with >70% coverage with >2 kb footprints. Dashed horizontal lines mark 10%, 5%, and 1% local false discovery rate (local FDR) thresholds, estimated from a comparison of frontal lobe and testis distributions across bins of the distribution. **E.** Histograms showing distribution of accessible patch sizes (n = 371,430) in deciles of >2 kb footprint occupancy in testes with color gradient indicating correspondence with Figure 2C. Lacuna size distribution of sperm is indicated at bottom in black. **F.** Subsamples of reads (n = 100) in the first bin, last bin, and sperm, sorted by length of accessible patch. Orange, protamine footprints; green, nucleosomal footprints; blue, subnucleosomal footprints.

Quantifying individual testis Fiber-seq reads based on their density of >2kb footprints revealed that testis chromatin fibers originate from cells at diverse stages of histone-protamine exchange (**Fig. 2C**). Many reads were exclusively occupied by nucleosome-sized footprints, identical to the paired somatic tissue (brain frontal lobe, **Fig. 2B**), consistent with fully chromatinized spermatogenic precursors or testicular somatic cells (**Fig. 2C**). In contrast, >2kb footprints were largely absent from somatic tissue, supporting their selective marking of protamine occupancy. Binning testis reads by overall protamine occupancy showed that increasing protamination was associated with a progressive reduction in the nucleosome density within lacunae (**Fig. 2D–F, Supplemental Fig. S2B**). However, fewer than 10% of testis fibers fully matched the profile of mature sperm. Specifically, although protamine lacunae in testis were similar in length to those in sperm, these lacunae retained a higher density of nucleosomes in testis (**Fig. 2D**). These results suggest that protamine deposition is initially impeded at lacunae containing normally spaced nucleosomes, followed by progressive nucleosome eviction and increasing sparsity within these regions during subsequent stages of spermiogenesis.

### Determinants of protamination and nucleosome retention in sperm

We next investigated whether specific chromatin or genomic features modulate the nucleosome-to-protamine transition during spermatogenesis (**Supplemental Fig. S2C-D**). Overall, in sperm we observed a modest enrichment of both protamine lacunae and retained nucleosomes at transcription start sites (TSS) of protein-coding genes and tRNA genes (**Fig. 3A, Supplemental Fig. S2E**), with protamine lacunae being preferentially enriched at gene promoters associated with developmental functions in either germ or somatic cells (**Fig. 3B-E, Supplemental Fig. S2F, Supplemental Table S1**). Incorporating testis RNA-seq data revealed that promoters for genes highly expressed in testis are strongly enriched for harboring protamine lacunae in sperm (**Fig. 3H**), whereas enrichment for retained nucleosomes in sperm was similar for high- and low-expressed testis genes (**Fig. 3I**), consistent with retained paternal chromatin information being predominantly encoded in protamine lacunae compared to nucleosome retention in sperm. Genes highly expressed in testis already showed evidence of protamine depletion in testis chromatin, while retaining a stereotypical nucleosome phasing pattern relative to the TSS not seen for low-expressed genes (**Supplemental Fig. 2G-H**), as expected if these nucleosomes are retained at positions they occupied as a result of transcriptional activity in early spermiogenesis. Together, these findings indicate that high transcriptional activity during spermatogenesis causes discontinuities in protamination along with subsequent retained nucleosomes in sperm (**Fig. 3J**).

**Figure 3.**
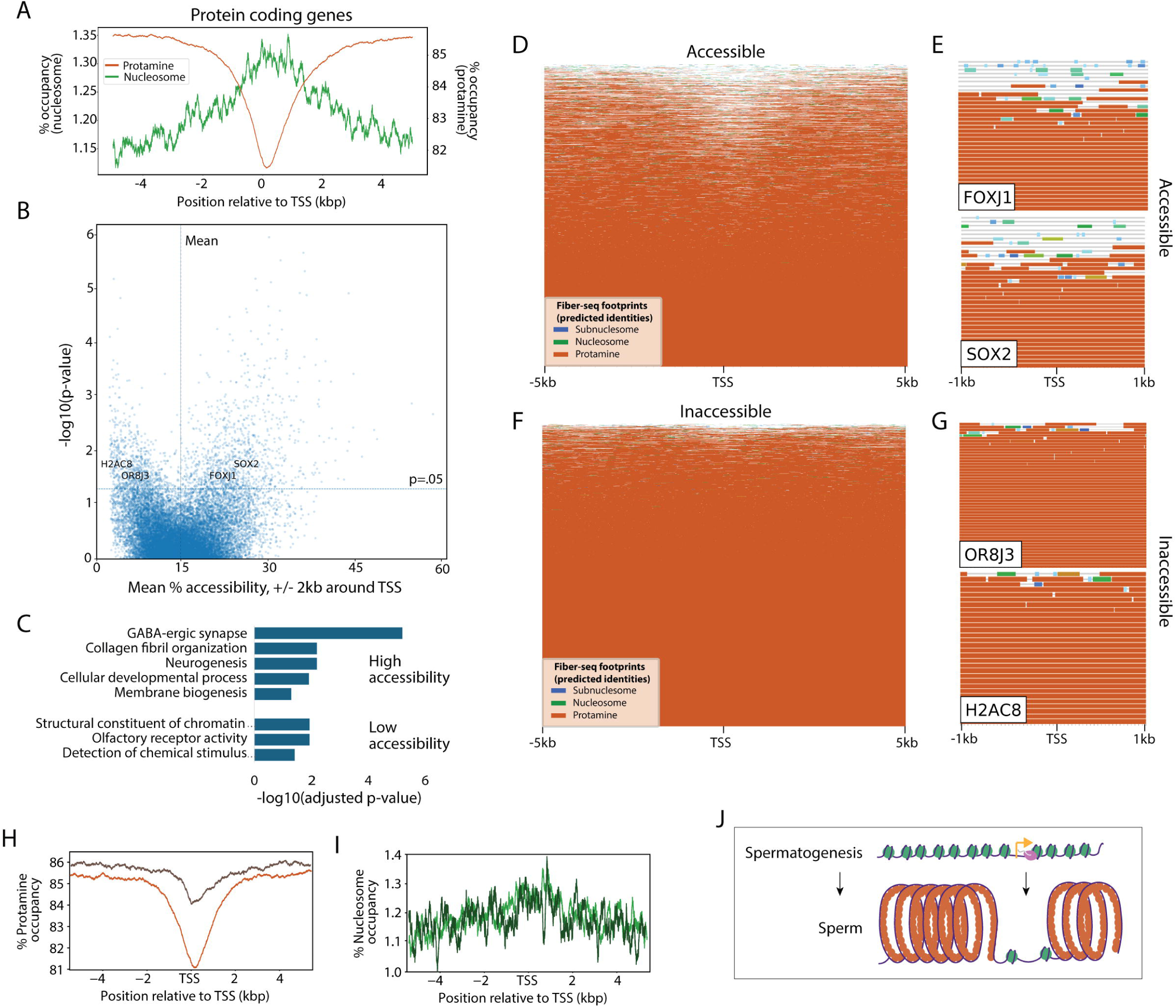
Preferential nucleosome retention and protamine depletion at transcriptionally active and developmental promoters. **A.** Metaplots of protamine and nucleosome occupancy within 5 kilobases (kb) upstream and downstream of transcription start sites for all protein coding genes. **B.** Identification of promoters with significantly greater mean accessibility than expected by chance. X-axis shows the mean accessibility of reads overlapping the promoter region (TSS +/− 2kb); y-axis shows p-value based on expected mean accessibility using a ranksums test. TSSs shown in panels E and G are labeled. Threshold lines indicate the mean value of the mean accessibility and p < .05. **C.** Gene ontology categories enriched among promoters with significantly high or low accessibility. **D.** Sample of highly accessible reads, centered on the TSS and sorted by length of accessible patch (n = 500). Orange, protamine footprints; green, nucleosomal footprints; blue, subnucleosomal footprints. **E.** Two examples of promoters in the high accessibility category. Reads are centered on the TSS and sorted by length of the accessible patch (n = 38, n = 46). Orange, protamine footprints; green, nucleosomal footprints; blue, subnucleosomal footprints. **F.** Sample of highly inaccessible reads, centered on the TSS and sorted by length of the accessible patch (n = 500). Orange, protamine footprints; green, nucleosomal footprints; blue, subnucleosomal footprints. **G.** Two examples of promoters in the low accessibility category. Reads are centered on the TSS and sorted by length of the accessible patch (n = 56, n = 32). Orange, protamine footprints; green, nucleosomal footprints; blue, subnucleosomal footprints. **H.** Metaplot of protamine occupancy in sperm chromatin fibers +/− 5kb from the promoter of the top (orange) or bottom (brown) quintile of expressed genes in testis (n = 10^6^). **I.** Metaplot of nucleosome occupancy in sperm chromatin fibers +/− 5kb from the promoter of the top (light green) or bottom quintile (dark green) of expressed genes in testis (n = 10^6^). **J.** Model for the effect of high transcription during late spermatogenesis on protamine and nucleosome profiles in sperm.

We next sought to understand if particular histone post-translational modifications (PTMs) play a role in modulating the protamine-to-nucleosome transition, as many PTMs are known to be present on histones retained in sperm ^5,29,30^. Using ChIP-seq data we generated from human sperm for the transcription-associated PTMs H3K27ac and H3K4me3 and for the repressive PTM H3K27me3, we observed that ChIP-seq defined sites (peaks) of retained nucleosomes were significantly and selectively enriched in nucleosome footprints and protamine lacunae in sperm Fiber-seq data (**Fig. 4A-C; Supplemental Table S2**), further confirming the accuracy of our Fiber-seq data for measuring nucleosome retention. We observed that loci containing H3K27ac histones or both H3K4me3 and H3K27me3 histones (*i.e.,* bivalent histones)^31–33^ exhibited the highest rate of nucleosome retention and protamine lacunae. Notably, unlike H3K27ac histones, which mark highly expressed genes, sites marked by bivalent histones are typically lowly expressed in testis ^34–37^ (**Fig. 4B-C**), but have previously been shown to be retained in sperm and enriched at elements critical for early development^5,6,12^. Together, our results indicate that protamine lacunae and retained nucleosomes are preferentially enriched at two classes of TSS in sperm: genes highly transcribed during late stages of sperm development, and genes transcriptionally repressed but marked by bivalent chromatin. Importantly, we find that most nucleosomes are evicted by the time sperm are fully mature, including those with PTMs, indicating that protamine lacunae are the dominant form of potential epigenetic inheritance at these loci in sperm (**Fig. 3J, 4D, 4H**).

**Figure 4.**
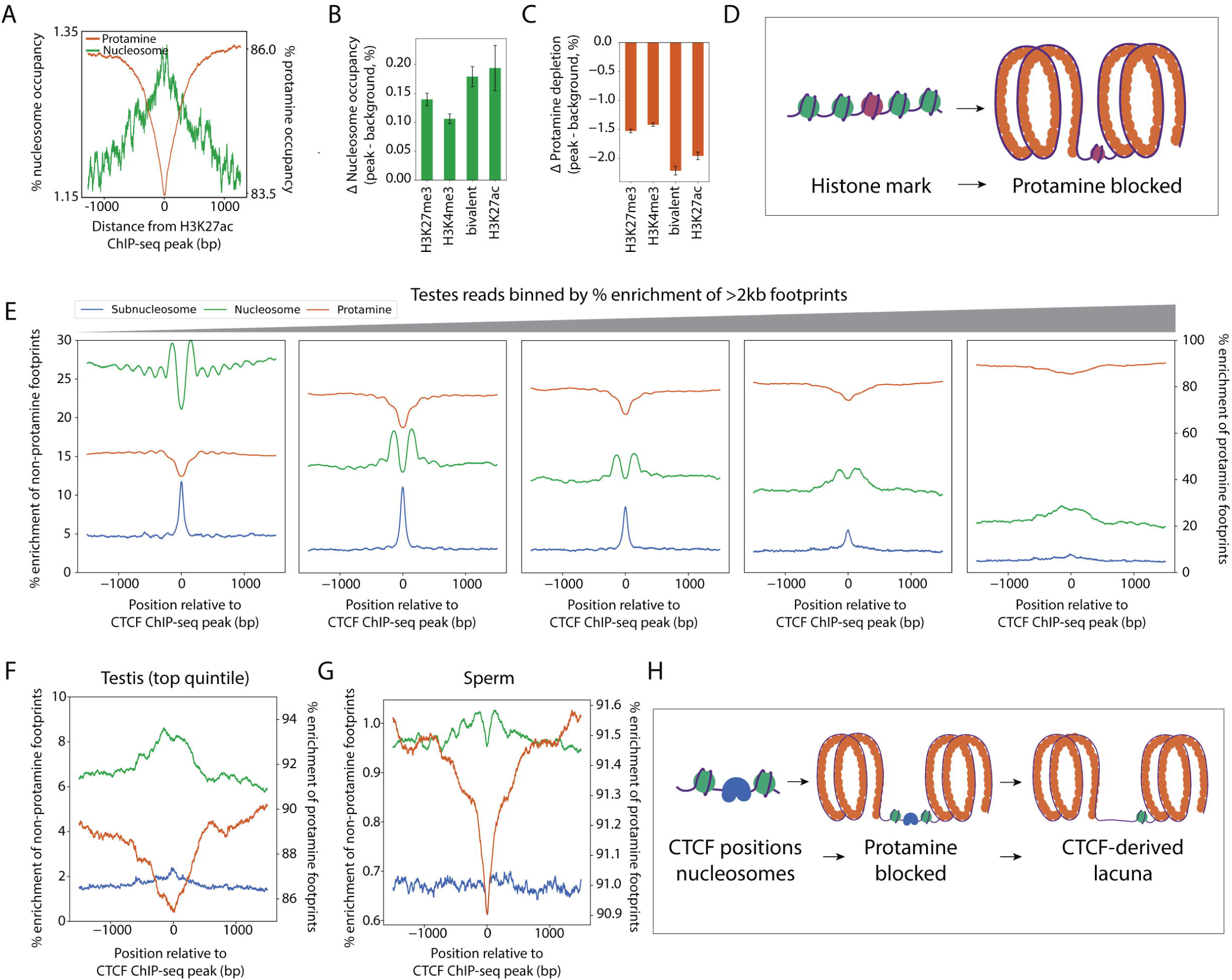
Histone marks and CTCF influence patterns of protamine lacunae. **A.** Metaplots of protamine and nucleosome occupancy centered on H3K27ac peaks in sperm (n = 452,871). **B.** Barplot showing the increase in percent nucleosome occupancy over background for reads centered on ChIP-seq peaks corresponding to bivalent, H3K27ac, H3K4me3, or H3K27me3 histone PTMs (n = 3,508,699, n = 452,871, n = 9,116,193, n = 4,031,007). Error bars are derived from two standard deviations from the background distribution of nucleosome footprint occupancy 4-5kb from the peak. **C.** Barplot showing the decrease in percent protamine occupancy over background for reads centered on ChIP-seq peaks corresponding to bivalent, H3K27ac, H3K4me3, or H3K27me3 histone PTMs (n = 3,508,699, n = 452,871, n = 9,116,193, n = 4,031,007). Error bars are derived from two standard deviations from the background distribution of protamine footprint occupancy 4-5kb from the peak. **D.** Model for the effect of histone modifications during spermatogenesis on protamine profiles in sperm. **E.** Metaplots of subnucleosomal (<90bp), nucleosome (90-200bp), and protamine (>200bp) percent occupancy on testis chromatin fibers binned into quintiles of >2kb footprint occupancy (n = 321,505 reads per quintile), centered on whole-testis ChIP-seq peaks for CTCF. **F.** Zoomed metaplot of subnucleosomal (<90bp), nucleosome (90-200bp), and protamine (>200bp) percent occupancy on reads from the highest quintile of testis >2kb gene expression (n = 321,505), centered on whole-testis ChIP-seq peaks for CTCF. **G.** Metaplot of subnucleosomal (<90bp), nucleosome (90-200bp), and protamine (>200bp) percent occupancy on reads from sperm (n = 5,089,124), centered on whole-testis ChIP-seq peaks for CTCF. **H.** Model for the effect of transcription factor binding during spermatogenesis on protamine profiles in sperm.

Finally, we sought to evaluate whether transcription factor (TF) occupancy modulates the nucleosome-to-protamine transition. Using ChIP-seq data from human testis, we observed that numerous protein-bound DNA binding elements in testis, including those bound by CTCF, were focally enriched in protamine lacunae in sperm (**Fig. 4G, Supplemental Fig. S3**), indicating that TF occupancy at these sites in testis is associated with alterations in the nucleosome-to-protamine transition. However, although CTCF has been reported to be present on sperm chromatin^13,16,17^, we did not observe any evidence of subnucleosomal footprints at CTCF binding elements in sperm, indicating that the effect of CTCF occupancy on protamine lacunae persists beyond the time period when CTCF is physically present at those elements. To explore this further, we evaluated occupancy of subnucleosomal footprints, nucleosomal footprints, and larger protamine footprints along testis fibers that were binned into quintiles based on their overall levels of >2kb protamine footprints (**Fig. 2D**). This revealed that protamine lacunae form at CTCF-bound elements at early stages of protamine deposition, and that late stages of the nucleosome-to-protamine transition are associated with loss of subnucleosomal footprints but retention of nucleosomal footprints and protamine lacunae at these sites, indicating that CTCF occupancy in the testis guides this chromatin effect in sperm (**Fig. 4E, F**). Together, these findings indicate that sperm retain a chromatin memory of DNA binding for certain TFs in the form of protamine lacunae, even though the proteins themselves are not retained (**Fig. 4H**), analogous to the effect observed for late-evicted nucleosomes.

### Centromere kinetochore binding regions selectively retain nucleosomes in sperm

In addition to histone PTMs, the histone H3 variant CENP-A has been shown to be abundant in sperm, where it can transmit information about centromere specification to the embryo^9–11,38^. Recent telomere-to-telomere (T2T) studies have revealed that CENP-A occupies only focal 20-100 kb regions of each chromosome’s α-satellite higher order repeat (HOR) (*i.e.,* the kinetochore binding region), which is marked by a characteristic pattern of hypo-CpG methylation (*i.e.,* the centromere dip region [CDR])^39,40^. Furthermore, the location of a CDR within a chromosome’s α-satellite HOR is largely epigenetically stable from parent to offspring^41^.

To resolve whether the focal retention of CENP-A histones within CDRs could mediate intergenerational inheritance of the precise location of the kinetochore binding region, we applied sperm Fiber-seq to cryopreserved sperm collected from the individual who is the genetic basis for the HG002 genome assembly, which includes accurate assemblies of all 46 centromeres^42^. We observed that the extent of nucleosome-to-protamine transition differed between various classes of repeat elements (**Supplemental Fig. S4A-D**), with α-satellites showing amongst the highest rate of nucleosome retention genome-wide (**Fig. 5A**). However, nucleosome retention within α-satellites appeared to be focally enriched at CDR positions as defined using somatic tissue from HG002 (**Fig. 5B-C**).

**Figure 5.**
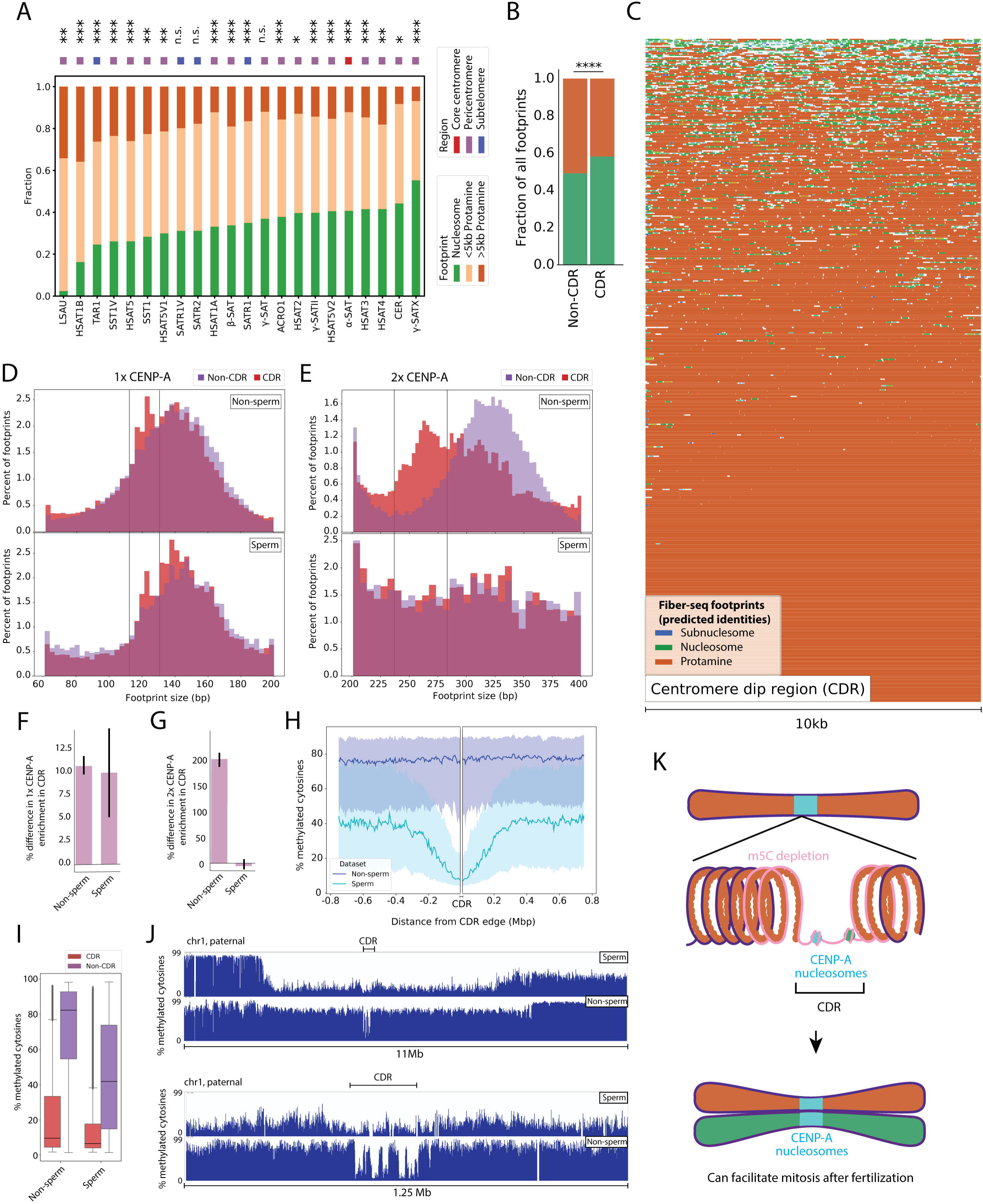
High CENP-A nucleosome retention in centromere dip regions. **A.** Fraction of total nucleosomal, <5kb protamine, or >5kb protamine footprints in reads containing each class of satellite sequence (counts available in Table SX). Designation of each satellite as associated with subtelomeric, pericentromeric, and centromeric regions is indicated by blue, purple, and red squares. Significance is based on a chi-squared test of the footprint distributions compared to a randomly sampled background of non-satellite sequences of equal length with Benjamini-Hochberg multiple testing correction (n.s.: p>.05, *p<.05, **p<10^2^, ***p<10^3^). **B.** Enrichment of total counts of individual nucleosome and protamine footprints (<200bp, >200bp, n = 80,870) in sampled reads (n = 3,142) from centromeric non-CDR regions compared to CDR regions. Significance is indicated above plot (chi-squared test, ****p < 10^−144^, n = 3,142). **C.** Individual Fiber-seq reads from the centromere dip region (CDR) in HG002 sperm sorted from low to high protamine content (n = 500). Orange, protamine footprints; green, nucleosomal footprints; blue, subnucleosomal footprints. **D.** Distribution of mononucleosome footprint sizes from reads within CDR (red) and sampled non-CDR (purple) centromere regions (n = 3,142) in data from sperm and non-sperm (lymphoblastoid cell line) samples. The size range for 1x CENP-A footprints is indicated with lines overlaid on the plot. **E.** Distribution of dinucleosome footprint sizes from reads within CDR (red) and sampled non-CDR (purple) centromere regions (n = 3,142) in data from sperm and non-sperm (lymphoblastoid cell line) samples. The size range for 2x CENP-A footprints is indicated with lines overlaid on the plot. **F.** Barplot showing quantification of the percent enrichment of 1x CENP-A footprints in CDR vs non-CDR regions (non-sperm: n = 12,118, sperm: n = 3,667). Errorbar indicates a 95% confidence interval from 1000x bootstrapped resamplings. **G.** Barplot showing quantification of the percent enrichment of 2x CENP-A footprints in CDR vs non-CDR regions (non-sperm: n = 147,295, sperm: n = 1,516). Errorbar indicates a 95% confidence interval from 1000x bootstrapped resamplings. **H.** Plot of median % m5C methylation starting at the edge of all CDRs in sperm and non-sperm datasets. CDRs located within 500kb of each other are merged. Error bars represent the 25th to 75th percentile interval. **I.** Boxplot showing distribution of CpG methylation rates for non-CDR and CDR centromere regions in non-sperm (lymphoblastoid cell line) and sperm datasets. **J.** CpG methylation (5mC) at the CDR and surrounding centromeric region in sperm and non-sperm (lymphoblastoid cell line) samples. CDR is indicated. Tracks are shown at two different scales in the top (11 Mb) and bottom (1.25 Mb). **K.** Model showing the relative extent of CENP-A containing nucleosomes (teal) and hypomethylated DNA (pink). CENP-A nucleosomes mark the CDR, forming the site of future kinetochore assembly to facilitate mitosis after fertilization.

We have previously found that CENP-A-containing nucleosomes within CDRs from somatic tissues produce characteristic Fiber-seq footprints corresponding to CENP-A-containing mono-nucleosomes (∼120bp), and CENP-A-containing di-nucleosomes (∼260bp) ^43^. Similarly, we observed a strong enrichment of ∼120bp footprints selectively within CDRs (**Fig. 5D,F**), indicating that CENP-A-containing mono-nucleosomes are focally retained at somatic CDRs within sperm. Notably, the relative proportion of CENP-A-containing mono-nucleosome footprints to canonical (∼146bp) mono-nucleosome footprints was similar between sperm and somatic cells, suggesting that these nucleosomes are strongly retained during the nucleosome-to-protamine transition (**Fig 5D)**. In contrast, we did not observe retention of footprints corresponding to CENP-A-containing di-nucleosomes (**Fig. 5E,G**), consistent with these di-nucleosomes corresponding to sites of kinetochore attachment in somatic cells ^43^, as there are no kinetochores present in sperm. Notably however, despite the focal retention of nucleosomes and CENP-A-containing mono-nucleosomes at somatic CDRs within sperm, the characteristic pattern of focal hypo-CpG methylation at these somatic CDRs was not retained in sperm. Rather, in sperm the entire α-satellite HOR appeared to display an overall reduction in CpG methylation, with the focal area of hypo-CpG methylation spreading by several hundred kilobases surrounding the CDR (**Fig. 5H-J**). Together, these results indicate that the focal retention of CENP-A mono-nucleosomes, rather than DNA hypomethylation, marks the CDR and thus may play a central role in retaining centromere-defining epigenetic information in the paternal gamete (**Fig. 5K**).

### Sub-motile sperm display altered nucleosome-to-protamine landscapes

Having established a genome-wide map of nucleosome-to-protamine transition, we next sought to resolve how disruptions in this process might impact human health. As defects in nucleosome-protamine replacement have been linked to impaired sperm function in the form of immotility^44–50^, we sought to leverage sperm Fiber-seq to resolve the specific defects that may be mediating this process. Application of Fiber-seq to severely asthenospermic samples (*i.e.,* <15% motile sperm^51^) and normospermic controls (*i.e.,* >42% motile sperm) revealed a strong relationship between protamine content detected by Fiber-seq and fraction of motile sperm (**Fig. 6A**, **Supplemental Fig. S5A-D**). Specifically, lower motility was associated with higher protamine content (R=-.832, p=.011 with outlier, R=-.907, p=.005 without outlier; **Fig. 6B**), contrasting with the prevailing assumption that more protamine loading correlates monotonically with better sperm function in mammals^47,52,53^. In addition, sperm motility was also associated with nucleosome retention levels (R=.481, p=.228 with outlier, R=.895, p=.006 without outlier; **Fig 6C**), with most asthenospermic samples having sparse nucleosome retention paired with high protamine occupancy levels. In contrast, one of the asthenospermic samples showed high nucleosome retention paired with high protamine occupancy levels (**Fig. 6C**), a pattern mirroring that of immature stages of spermatogenesis as observed in the testis Fiber-seq data (**Fig. 2D-F**). This further indicates that protamination and nucleosome retention can occur independently of each other during spermatogenesis, and that Fiber-seq can accurately distinguish diverse aberrant mechanisms underlying functional sperm defects, even at very low sequencing coverage (**Supplemental Fig. S5A-D**).

**Figure 6.**
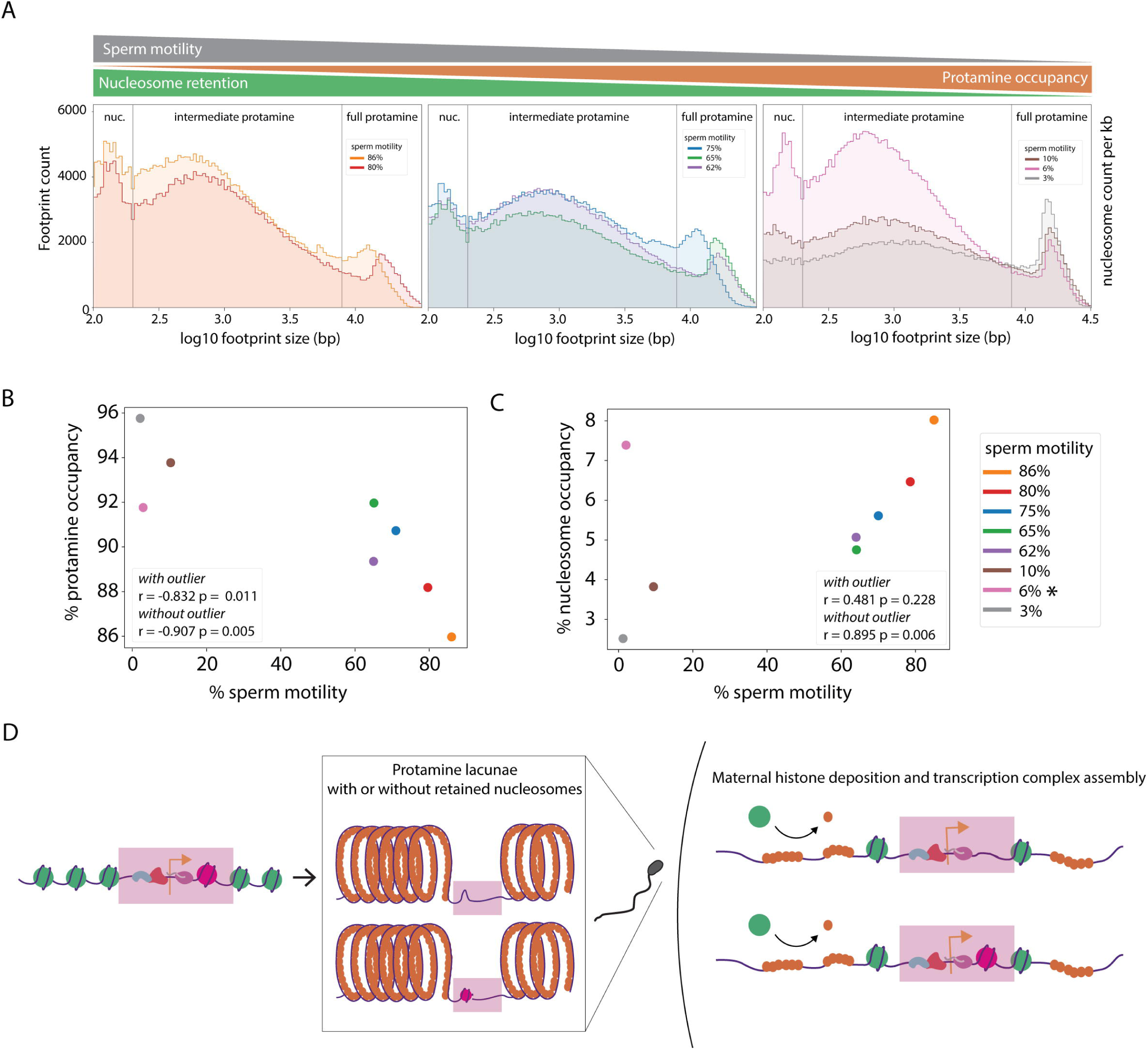
Impaired sperm motility is associated with increased protamine occupancy. **A.** Distribution of footprint sizes in sperm samples with high (left), intermediate (middle), or poor motility (right). Grey triangle above the plots indicates motility; orange and green triangles indicate inferred levels of protamine and nucleosome occupancy, respectively (reads for each sample sampled to 79Gb each). **B.** Protamine occupancy vs. percentage of motile sperm for the same samples (sampled to 79Gb each) shown in A. Outlier data point (pink) is marked by an asterisk in the key. Pearson correlation p and r values with and without the outlier are indicated on the plot. **C.** Nucleosome occupancy vs. percentage of motile sperm for the same samples (sampled to 79Gb each) shown in A. Outlier data point (pink) is marked by an asterisk in the key. Pearson correlation p and r values with and without the outlier are indicated on the plot. **D.** Proposed model where transcriptional activity in late spermiogenesis results in protamine lacunae with or without retained nucleosomes in mature sperm. Either persistent protamine depletion and/or associated nucleosome retention biases chromatin accessibility and may alter the dynamics of remodeling and genome activation after fertilization.

## Discussion

Together, our results illuminate the specific architecture by which epigenetic information encoded in chromatin is retained in sperm. Notably, the retention of this information is probabilistic rather than deterministic, as no locus demonstrated the complete focal retention of epigenetic information in all sperm, arguing against a model where programmed histone retention is essential for function of specific genes after fertilization^54^. However, focal retention of epigenetic information in sperm is not random, as specific genomic regions, including some with critical functional roles, are much more likely than others to retain epigenetic information in sperm chromatin. Importantly, epigenetic information in sperm is predominantly encoded in the form of protamine lacunae: discontinuities in protamination of the DNA fiber that selectively localize to sites of active transcription at the time of protamine loading, occupancy sites for certain TFs, and post-translationally modified histones. Nucleosomes are also preferentially retained in some genomic contexts. Specifically, these include CENP-A-containing mono-nucleosomes at the kinetochore binding region of the centromere, well-positioned nucleosomes at TSSs of genes with high transcriptional activity during the process of protamination, and nucleosomes containing specific post-translationally modified histones, such as bivalent H3K27me3/H3K4me3 nucleosomes.

Our data is consistent with a model whereby expanding blocks of protamine-coated DNA are disrupted by specific chromatin features during protamine exchange, including nucleosomes and TFs. These discontinuities in the protamine fiber are initially populated by retained nucleosome arrays, which are then largely evicted during the final stages of spermatogenesis, leaving protamine lacunae that contain large stretches of unpackaged DNA with scattered occupancy of retained mono- or di-nucleosomes. This model suggests that nucleosomes of differing “fragility”^13,55,56^ differentially impact the process of protamination such that the main outcome is the creation of protamine lacunae in sperm at sites of more stable nucleosomes, while the nucleosomes themselves may or may not be retained (**Fig. 6D**). Consequently, the process of nucleosome-to-protamine exchange leaves a clearing of accessible DNA at critical regulatory elements, potentially biasing activation of these sites after fertilization. However, we find that this process can be decoupled, wherein sperm with robust protamination but inadequate nucleosome eviction are associated with male-factor infertility.

Although protamine lacunae appear to be the dominant manner in which the paternal epigenome is retained genome-wide in sperm, the centromere appears to rely more heavily on nucleosome retention for the focal transmission of the kinetochore binding region. Specifically, locations that are more enriched for CENP-A in somatic tissues preferentially retain CENP-A-containing mono-nucleosomes in sperm, despite sperm lacking the characteristic focal hypo-CpG methylation pattern at these sites. This suggests that focal transgenerational inheritance of the centromere location from the paternal lineage is more likely to be encoded by retained CENP-A nucleosomes as opposed to differential CpG methylation.

Finally, our data hold significant potential for clinical testing in male infertility. Defects in sperm histone/protamine balance have been associated with abnormal semen parameters, unexplained failure of assisted reproductive technologies, and infertility. While the motility defects assayed here are easily detectable by standard semen analysis, we found that Fiber-seq revealed previously undetected differences in protamine and nucleosome density among low-motility samples. Fiber-seq may have the potential to reveal underlying functional defects that would be hidden using standard clinical approaches and is highly scalable, suggesting that it may be efficient and cost-effective in diagnostic applications.

## Acknowledgments

We thank Michael Levine and L. Stirling Churchman for feedback and Zuri Omari-Muhammad for discussions. A.B.S. and B.J.L. are Pew Scholars, supported by the Pew Charitable Trust. PacBio HiFi sequencing was performed at the NIH Intramural Sequencing Center (NISC) as well as the UW PacBio sequencing core and the Yale Center for Genome Analysis. This work was supported by a grant from the American Society for Reproductive Medicine to B.J.L. and by NIH grants 1DP5OD029630 and 1U01HG013744 to A.B.S.. T.W.T. was supported by an NIH grant (GM118147) to Michael Levine at Princeton University. R.A.H. was supported by training grant 5T32HD007149-45 and S.C.B. was supported by training grant 2T32GM007454-46 from the NIH. This research is supported by the NIH Common Fund, through the Office of Strategic Coordination/Office of the NIH Director under awards U24 MH133204, U24 NS132103, UG3 NS132024, UG3 NS132061, UG3 NS132084, UG3 NS132105, UG3 NS132127, UG3 NS132128, UG3 NS132132, UG3 NS132134, UG3 NS132135, UG3 NS132136, UG3 NS132138, UG3 NS132139, UG3 NS132144, UG3 NS132146, UM1 DA058219, UM1 DA058220, UM1 DA058229, UM1 DA058230, UM1 DA058235, and UM1 DA058236. This work was supported, in part, by the Intramural Research Program of the US National Human Genome Research Institute, National Institutes of Health. The contributions of NIH authors are considered Works of the United States Government. The findings and conclusions presented in this paper are those of the authors and do not necessarily reflect the views of the NIH or the U.S. Department of Health and Human Services.

## Data availability

For the SMaHT donor tissue, all protected (e.g., sequencing data, germline variant calls, and complete donor metadata for the SMaHT Production donors) and open (e.g., gene expression quantification, tables, etc.) access data are available in dbGaP (phs004193 for the SMaHT Benchmarking data; phs004104 for the SMaHT Production data. The SMaHT Data can be accessed from the SMaHT Data Portal at https://data.smaht.org/.

## Author contributions

Conceptualization: TWT, RAH, ABS, BJL

Investigation: TWT, RAH, JR, SCB, DD, BM

Resources: CEM, AA, ES, AMP

Validation: TWT, RAH

Formal analysis: TWT, RAH

Visualization: TWT, RAH, ABS, BJL

Funding acquisition: ABS, BJL Supervision: ABS, BJL

Writing - original draft: TWT, RAH, ABS, BJL

Writing - review and editing: TWT, RAH, ABS, BJL

## Methods

### Human samples

De-identified and discarded human semen samples were obtained from the Yale Fertility Center and this work was determined to be non-human subject research under Yale IRB #2000032744.

### Sperm sample collection

To purify spermatozoa, samples were gently pipetted over a density gradient composed of 1 mL of 45% ORIGIO® Gradient on top of 1 mL of 90% ORIGIO® Gradient (#84010060). Gradients were spun at 400xg for 20 minutes to pellet out sperm, which were resuspended in ORIGIO® Sperm Wash (#84055060). Sperm count and motility were quantified via computer-assisted sperm analysis. Purified sperm were washed in PBS and treated with DNase (Qiagen #79254) to remove any extracellular DNA fragments. Sperm were then frozen in PBS at −80°C until use.

### Reconstituted DNA/protamine Fiber-seq

High molecular weight DNA was isolated from HEK 293 cells using the Promega Wizard® HMW DNA Extraction Kit (#A2920). The resulting DNA was treated with different concentrations of native human protamines (equal parts protamine 1 and protamine 2, Briar Patch Biosciences) for one hour at room temperature. All samples were then treated with 1 µL of the Hia5 methyltransferase for 10 minutes at 25°C before the addition of 3 µL of 20% sodium dodecyl sulfate (SDS, Teknova #S0294) to stop the methyltransferase reaction. Samples were mixed with 5 µL of Proteinase K (Promega # MC500C) and incubated for 15 minutes at 56°C before placement on ice for 1 minute. DNA was purified by adding 100 µL of phenol:chloroform:isoamyl alcohol (Sigma-Aldrich #P2069) followed by a short vortex and centrifugation at 16,100 x g for 10 minutes. The upper aqueous phase was transferred to new microfuge tubes, where DNA was precipitated with the addition of 4 µL of 5M NaCl and 800 µL of 100% ethanol. Samples were inverted to mix and incubated at −20°C for 30 minutes before centrifugation for 10 minutes at 16,100 x g followed by a wash with 200 µL of 70% ethanol. After removing supernatant, samples were inverted on paper towels to air dry for 15 minutes. DNA was reconstituted by rotation overnight in 100 µL of 10mM Tris-HCl (AmericanBio #AB14043) at room temperature and stored at 4°C for future use.

### m6A dot blot

Reconstituted DNA samples purified above were sonicated for 10 minutes in a Diagenode Bioruptor® at 4°C (30 seconds on/30 seconds off, 10 cycles). Each sample was concentration adjusted to 2 ng/µL and serially diluted. A PVDF membrane (Bio-Rad #1620177) was activated for 30 seconds in methanol and washed first in water and then in TBST (PBS with 0.25% Triton-X-100 and 0.1% Tween-20) before draining onto filter paper. The membrane was dotted with 2 µL of each sample and its serial dilutions, and then allowed to dry for 1.5 hours before blocking in 5% omniblock (AmericanBio #10109) diluted in TBST for 1 hour. The membrane was washed three times for 10 minutes with TBST and subsequently incubated for one hour in rabbit anti-N6-methyladenosine antibody (Sigma-Aldrich #ABE572) diluted 1:10,000 in blocking solution. The membrane was washed three more times in TBST and incubated for one hour in goat anti-rabbit IgG peroxidase antibody (VectorLabs #PI-1000) diluted 1:10,000 in blocking solution. After three final TBST washes, the membrane was incubated for 5 minutes in SuperSignal™ West Pico PLUS (Thermo Scientific #34580) and imaged on a ProteinSimple FluorChem E System.

### Sperm Fiber-seq

Frozen sperm samples of 10 million sperm each were thawed at room temperature, washed once in PBS, and resuspended in 1 mL PBS. Samples were treated with 1M dithiothreitol (DTT Thermo Scientific #20290) for two hours at room temperature, and then with 1M N-Ethylmaleimide (Sigma-Aldrich #E3876) for 30 minutes. Samples were then spun at 2,000 x g, washed in PBS, then spun again and resuspended in 60 µL of Buffer A (15mM Tris-HCl, 15mM NaCl, 60mM KCl, 1mM EDTA, 0.5mM EGTA, and 0.5mM spermidine in ultrapure water). Samples were mixed with 60 µL of Lysis Buffer (Buffer A with 0.05% IGEPAL, Sigma #I8896) and incubated on ice for 10 minutes. Tubes were then centrifuged at 2000 x g for 5 minutes and samples were resuspended in 57.5 µL Buffer A. To this was added 1.5 µL of 32mM S-adenosylmethionine (New England Biolabs #B9003S) and 1 uL of the Hia5 methyltransferase. The methyltransferase reaction was allowed to proceed for 10 minutes at 25°C, and was halted by the addition of 3 µL of 20% SDS and 1 µL of 100mM DTT. Samples were then mixed with 37 µL PBS, 500 µL HMW Lysis Buffer A (Promega #A294A), and 3 µL RNase A (Promega #A797A), inverted 5x to mix, and incubated at 37°C for 15 minutes. After this incubation, 20 µL of Proteinase K (Promega #MC500C) was added; tubes were inverted 10x to mix and incubated at 56°C for 10 minutes before placement on ice for 1 min. To this was added 200 µL of Protein Precipitation Solution (Promega #A795A), and samples were mixed 5x by pipetting vigorously with wide-bore pipettes. Samples were spun at 16,100 x g for 5 minutes, and the supernatant was poured into a new microfuge tube containing 600 µL of isopropanol to precipitate DNA. Tubes were inverted 16x to mix and spun for 2 minutes at 16,100 x g. The supernatant was removed and the DNA resuspended in 600 µL of 70% ethanol, with inversion 4x to mix. Samples were spun once more for 2 minutes at 16,100 x g. The supernatant was removed and the tubes allowed to dry inverted for 15 minutes. Tris-HCl (10mM) was added, and the DNA was allowed to reconstitute in this solution overnight before storage at −80°C. DNA was sequenced using the PacBio Revio instrument.

### Sperm ChIP-seq

Buffers were prepared as follows. Buffer 1 was composed of 15 mM Tris-HCL, 60 mM KCl, 5 mM MgCl2, and 0.1 mM EGTA in water. MNase buffer was composed of 85 mM Tris-HCl, 3 mM MgCl2, and 2 mM CaCl2 in water. Combined buffer was composed of equal parts Buffer 1 and MNase buffer with 0.3M sucrose. Wash buffer A was composed of 50 mM Tris-HCl, 10 mM EDTA, and 75 mM NaCl in water. Wash buffer B was composed of 50 mM Tris-HCl, 10 mM EDTA, and 125 mM NaCl in water. Elution buffer was composed of 0.1M NaHCO3, 0.2% SDS, 5 mM DTT, 10mM Tris-HCl, and 1mM EDTA in water.

Protein G Dynabeads™ (Thermo Fisher 10004D) were prepared by washing twice in 10mM Tris-HCl with 1mM EDTA. Beads were further washed twice combined buffer before incubation for three hours at 4°C with combined buffer mixed with 1mg/mL bovine serum albumin and 100uL of tRNA solution (Sigma Aldrich #R8508). Beads were then washed once with combined buffer, resuspended in combined buffer, and left at 4°C until further use.

Frozen sperm samples of 10-15 million sperm were thawed at room temperature, washed once with PBS, and resuspended in 300 µL of buffer 1 with 0.3M sucrose and 10mM DTT. To this was added 300 µL of buffer 1 with 0.3M sucrose, 10mM DTT, 0.5% TERGITOL™ NP-40 (Sigma Aldrich #NP40S), and 1% sodium deoxycholate (Sigma Aldrich #D6750). Samples were incubated on ice for 30 minutes and then split to 100 µL aliquots. To each aliquot was added 100 µL of MNase buffer with 0.3M sucrose and 400U/mL micrococcal nuclease S7 (Roche #10107921001). Each aliquot was incubated for exactly 5 minutes at 37°C before the reaction was quenched by adding 2 μL of 0.5 M EDTA. Aliquots were placed on ice for 5 minutes and centrifuged at 16,300 x g for 10 minutes. Supernatants of each individual sample were recombined, treated with 65 μL of 20x protease inhibitor (Roche #11836170001), and precleared by adding 50 μL of prepared beads and rotating at 4°C for one hour. Samples were then centrifuged and transferred to new microfuge tubes, with 100 μL of each sample reserved as input. 5 μg (H3K4me3, H3K27me3) or 1 μg (H3K27ac) of antibody were added (Abcam #ab8580 for H3K4me3, #ab6002 for H3K27me3, and #ab4729 for H3K27ac). Samples were then rotated overnight at 4°C. The next day, 50 μL of prepared beads were added, and samples were incubated rotating at 4°C for 6-8 hours before the antigen-bound beads were washed once in wash buffer A and twice in wash buffer B. Two elution steps were performed by adding elution buffer to the beads for 15 minutes at 65°C, and total elution volume was combined for each sample. Both immunoprecipitated chromatin and reserved input chromatin were incubated with 6 μL of RNase A (Sigma Aldrich #70856) for 30 minutes at 37°C, and then with 6 μL of proteinase K (Thermo Fisher # AM2548) overnight at 55°C. The next morning, DNA was isolated using the ChIP DNA Clean & Concentrator kit (Zymo #D5201). ChIP-seq libraries were prepared using either the KAPA HyperPrep Kit (Roche, KK8504) or the Watchmaker DNA Library Prep Kit (#7K0103-096).

### ChIP-seq data processing

Samples were sequenced on an Illumina NovaSeq instrument and multiplexed to a read depth of approximately 30 million reads per sample. Paired-end reads were filtered with FASTX-Toolkit v0.0.14, and orphan reads were discarded with SeqKit v2.3.1. Sequencing adapters were removed with cutadapt v3.4 and reads were aligned to the hg38 genome reference using Bowtie2 v2.4.2. Peaks were called using MACS2 v2.2.9.1. Bivalent regions were determined by intersecting H3K4me3 and H3K27me3 peaks for each sample using BEDTools v2.30.0. Final files of H3K4me+, H3K27me3+, H3K27ac+, and bivalent regions were determined by merging all peaks present in at least one sample.

### Fiber-seq processing

Fiber-seq reads were aligned with pbmm2 to the hg38 reference genome. m6As were called with fibertools-rs v0.6.2 using default parameters, and converted to bed format for downstream footprint calling. Only reads >10kb were included in downstream analyses.

### 5mC calling

5mC enrichment was called using pb-CpG-tools v3.0.0 with default parameters.

### FiberHMM

Footprints were called on Fiber-seq reads using FiberHMM v1.4. In brief, FiberHMM is based on a hidden Markov model with two hidden states– accessible and inaccessible. To account for sequence biases, at each position the model takes into account the +/−3bp sequence context with experimentally derived methylation rates from naked DNA and untreated control datasets generated from *Drosophila* S2 cells. Transition and starting probabilities were trained 20 times with initial probabilities picked from the Dirichlet distribution with all parameters set to 1, on 10,000 reads from sperm samples or somatic samples. The best of these two models were chosen and used for the corresponding samples.

### Defining footprint classes and lacunae

In general, subnucleosomal footprints were defined as less than 90bp footprints, nucleosome footprints were defined as footprints greater than 90bp and less than 200bp in length, and protamine footprints were defined as footprints greater than 200bp in length. For figures distinguishing smaller and larger protamine footprints, small protamine footprints were defined as 300-10000bp footprints and large protamine footprints were defined as greater than 10000 bp footprints. Lacunae were defined as gaps between protamine footprints greater than 100bp.

### Sampling reads

To allow for direct comparison between datasets with different levels of coverage (*e.g.* the athenospermic samples), reads were sampled such that an equal level of bp of coverage overall was represented. For specific regions (*e.g.* satellites, CDRs, centromeres), a random selection of regions not included in the query set in the genome were chosen to match the exact distribution of region sizes and used as the comparison. All samplings were repeated at least 10x to validate that conclusions held. For figures 1H-I, 10^6^ reads were selected at random to generate the distributions to maintain consistency.

### Boxplots of nucleosome density in testis

Reads from testis were split into ventiles based on fraction coverage of >2kb footprints. Similarly, the top 70% >2kb coverage frontal lobe reads were identified. Count of nucleosomes per kilobase within protamine lacunae > 600bp in each group of reads was calculated. A local false discovery rate (local FDR) for each bin of protamine fraction was calculated by comparing normalized histograms of the frontal lobe and testis density distributions. For each bin, the local FDR was defined as the proportion of the signal attributable to the frontal lobe relative to the combined frontal lobe and testis signal.

### Per-gene lacuna enrichment analysis

TSSs were based on the annotated start sites from NCBI RefSeq GCF_000001405.40-RS_2023_10. Reads were centered on each TSS, and the average accessibility (fraction of bp not found in a protamine footprint) for the +/− 2000bp region around the TSS was calculated for each of the reads. A p-value for the likelihood of observing this distribution of accessibility values by chance was generated via a ranksums test compared to the distribution of all reads. The mean accessibility was then plotted along with this p-value. Significantly accessible and inaccessible genes were then used for gene-set enrichment analysis.

### Histone ChIP-seq comparative nucleosome and lacunae enrichment

Reads were aligned to ChIP-seq peaks for histone marks in sperm within a 10kb window. The distribution of protamine enrichment and nucleosome enrichment were then calculated. The peak value was calculated as the top value within 1kb of the peak center. The background occupancy at peak regions was calculated as the mean value within the first 1kb and last 1kb of the window. A standard deviation was also calculated for the background based on the distribution of these values, with errorbars plotted representing 2 standard deviations of range.

### Comparison of counts of footprint classes

For satellites, segments of reads were filtered based on overlap with annotated satellites, with a randomized background sample selected from an identical distribution of non-satellite sequence lengths. Footprints were counted, and binned based on sizes into nucleosome (<10**2.5 bp), small protamines (<10**4 bp), or large protamines (>10**4 bp). The distribution of fraction footprints in each bins was compared with a chi-squared test between the background and satellite samples, with Benjamini-Hochberg multiple testing correction. In the case of CDRs and centromeres, the centromere reads were sampled to match the length of the CDR reads, and bins were simplified to >10**2.5 and <10**2.5. The two distributions were then directly compared with a chi-sequared test.

### Gene ontology analysis

Gene ontology analysis was performed in R using the GOStats package with annotation package org.Hs.eg.db ^57^.

### Subsampled correlation of athenospermic samples

Each subsampling state was resampled 1000x and pearson r and p were calculated, and a group was defined as significant if at least 90% of replicate p-values within that group were below 0.05. For groups passing this threshold, replicate p-values were combined using Fisher’s method to yield a single group-level p-value. Pearson r values were reported as a distribution.

### Confidence intervals for barplots

Unless otherwise noted, confidence interval errorbars for barplots were generated via 1000x bootstrapped resampling of the input data, using a confidence interval indicated in the figure legend.

### HG002 annotations

The HG002 sperm and LCL data was aligned to the HG002v1.1a assembly. Centromere and satellite annotations were derived from https://github.com/hloucks/CenSatData.

### Additional datasets

Transcription factor ChIP-seq datasets were obtained from Chip-atlas. Testis expression bins were derived from The Human Protein Atlas version24.0 consensus tissue RNA database.

### Plotting libraries

Plots were generated with matplotlib and seaborn. Statistical analyses were carried out with scipy and numpy. m5C genome browser snapshots were generated with IGV and footprint genome-browser snapshots were generated with a custom visualization script.

