## Supplementary figures and images for "Protamine lacunae preserve the paternal chromatin landscape in sperm"

### Supplemental Figures

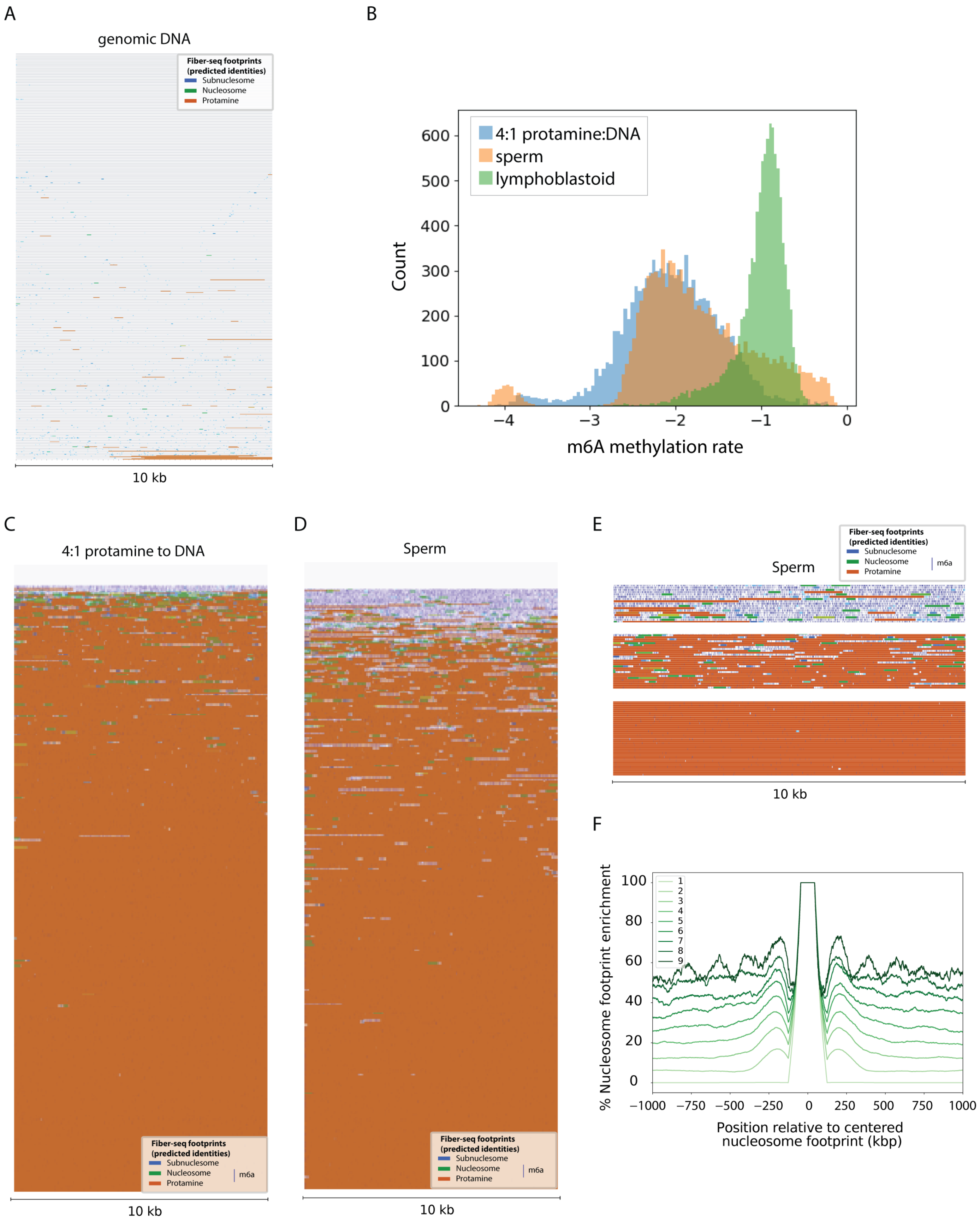

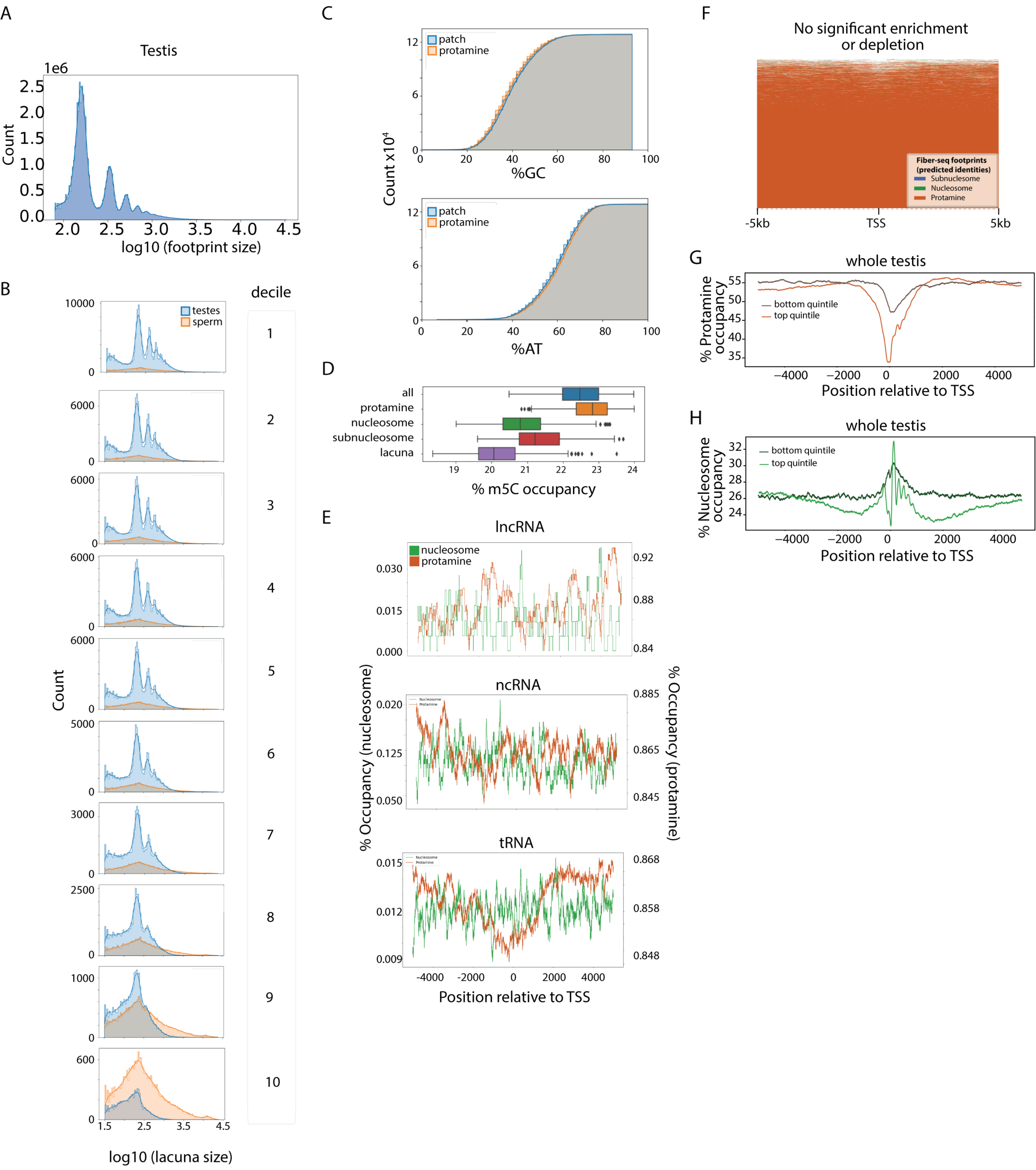

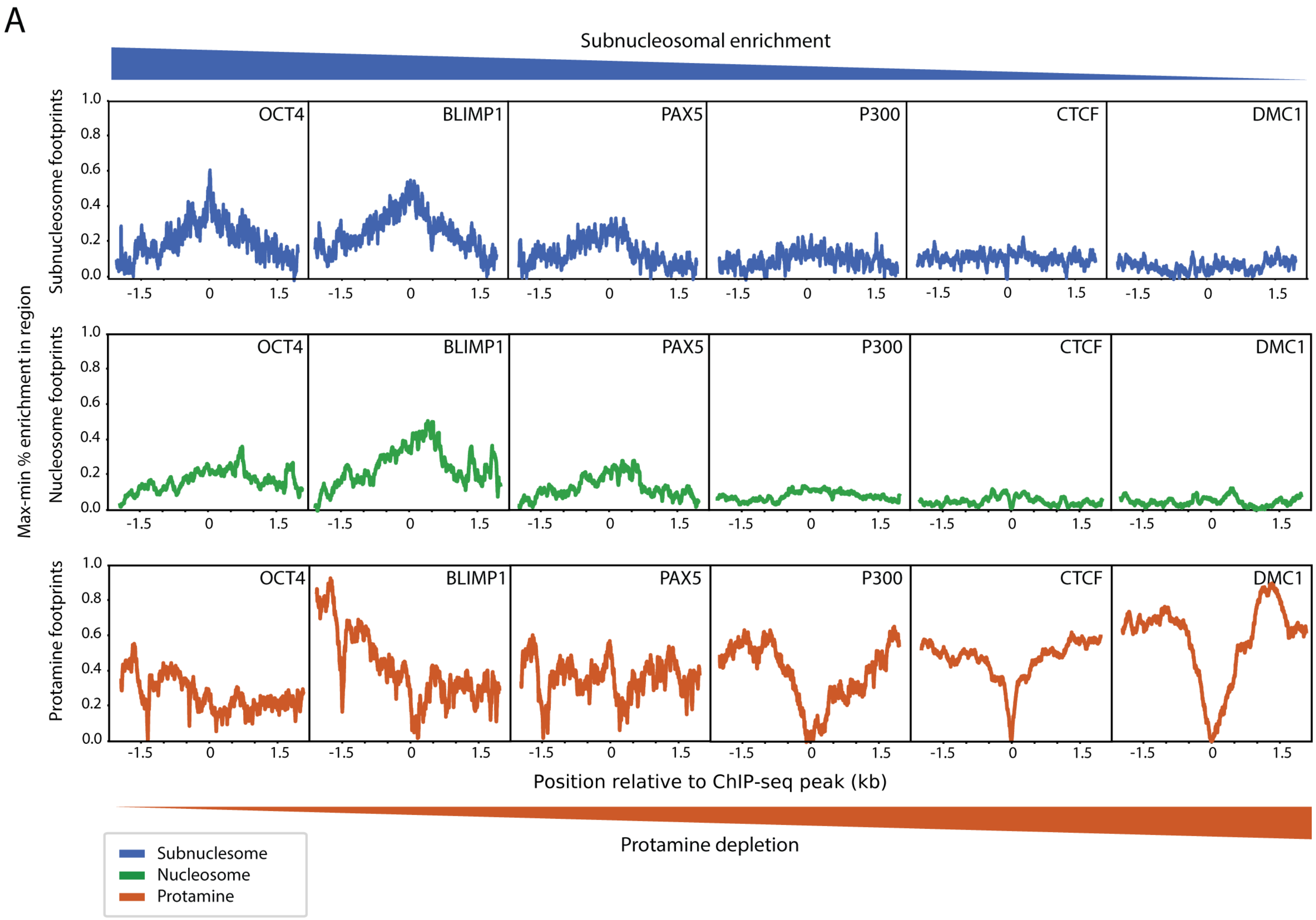

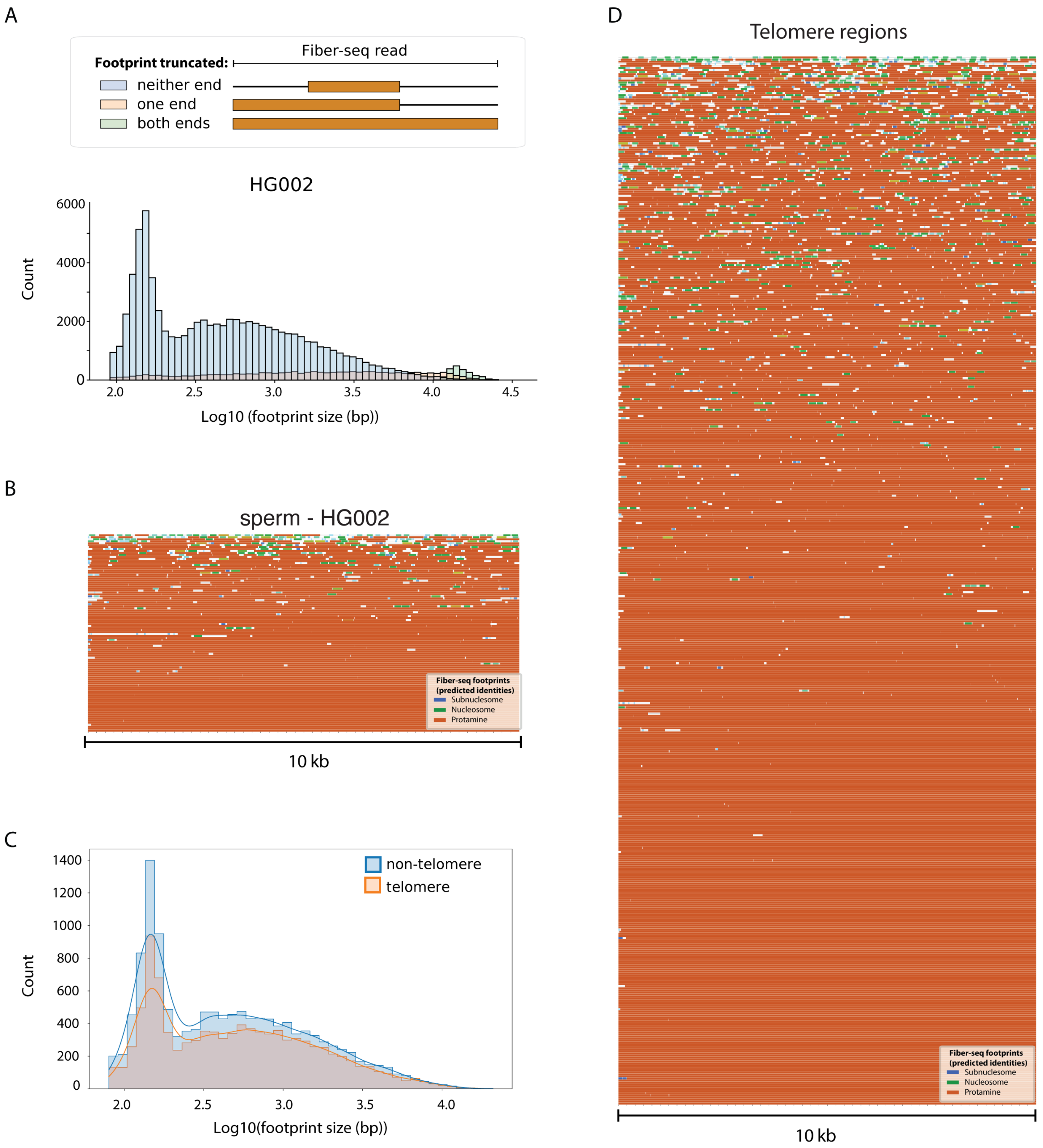

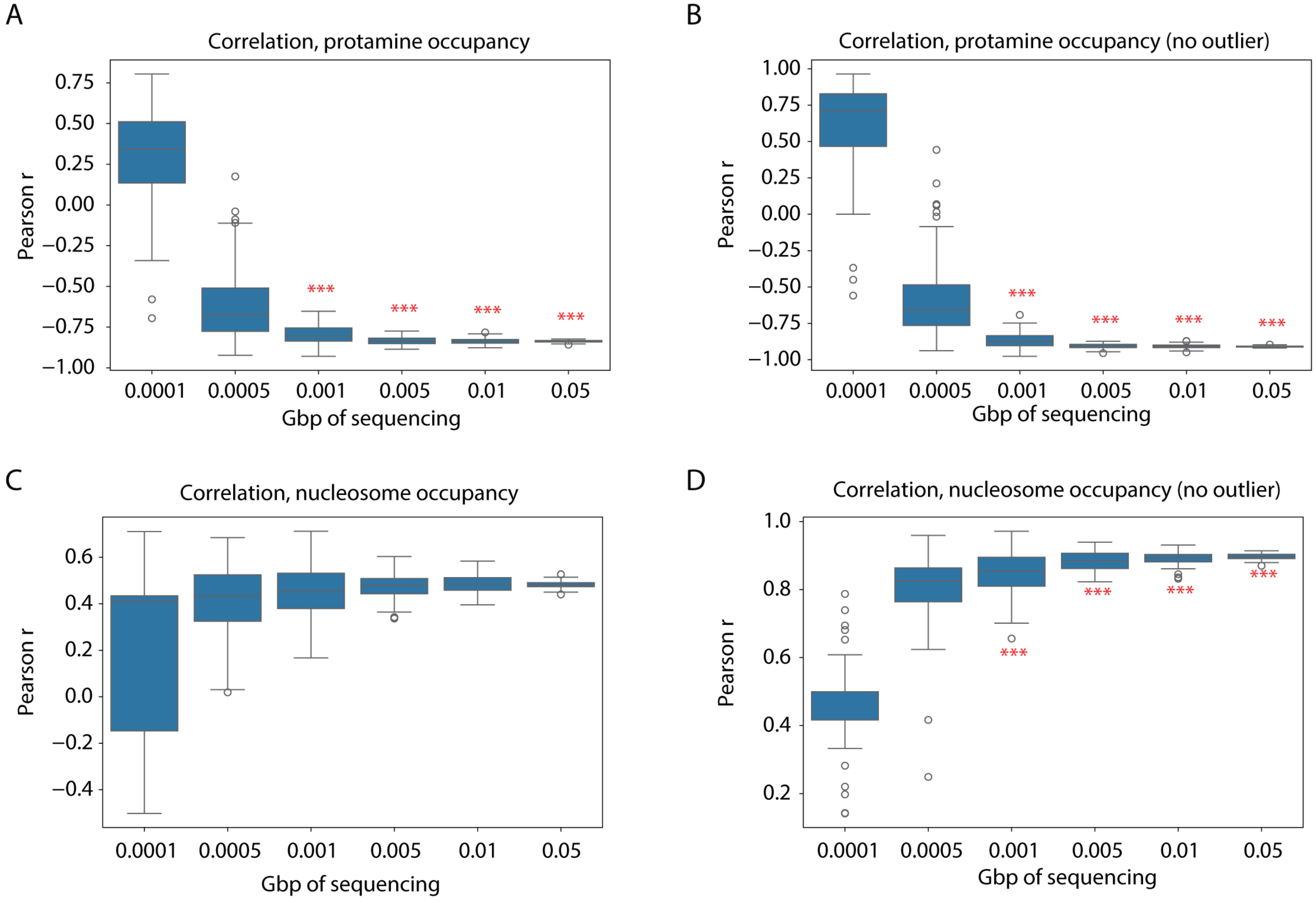
